## Supplementary material for "The Role of Experience in Prioritizing Hippocampal Replay": All supplementary figures

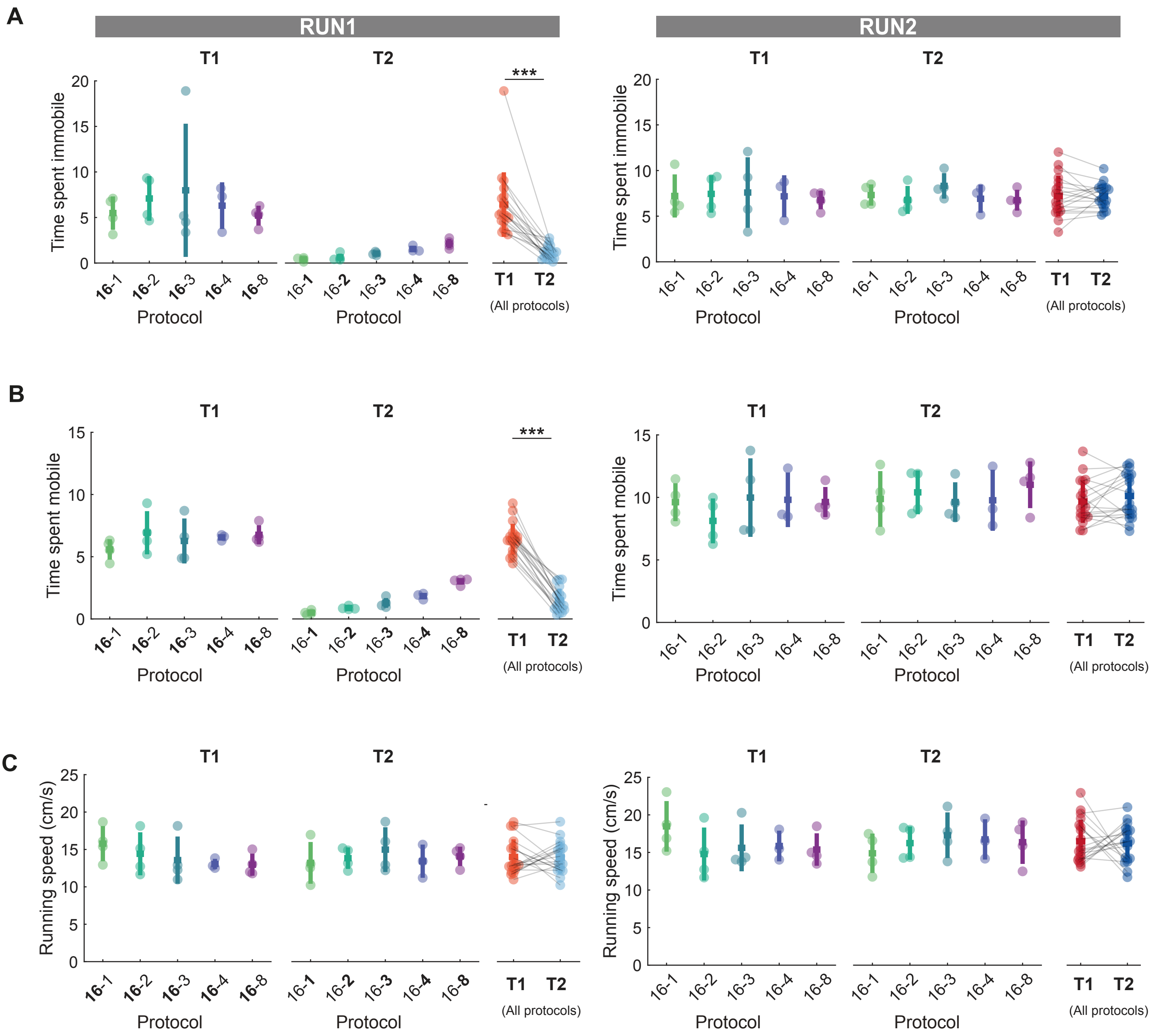


**Fig S1.** Summary of animal behavior during RUN1 and RUN2 for each protocol. **(A-C)** The behavioral metrics on track 1 and track 2 during RUN1 (left) and RUN2 (right) for each protocol. The lap number associated with each track during RUN1 was made in bold. The lap number was not highlighted as the time spent on two tracks were nearly identical. Each data point represents the mean theta sequence for a given track within a session, color-coded according to the experimental protocol and track identity (***p<0.001, two-tailed Wilcoxon signed rank test). **(A)** Time spent immobile for each protocol. **(B)** Time spent mobile for each protocol. **(C)** Moving speed for each protocol.

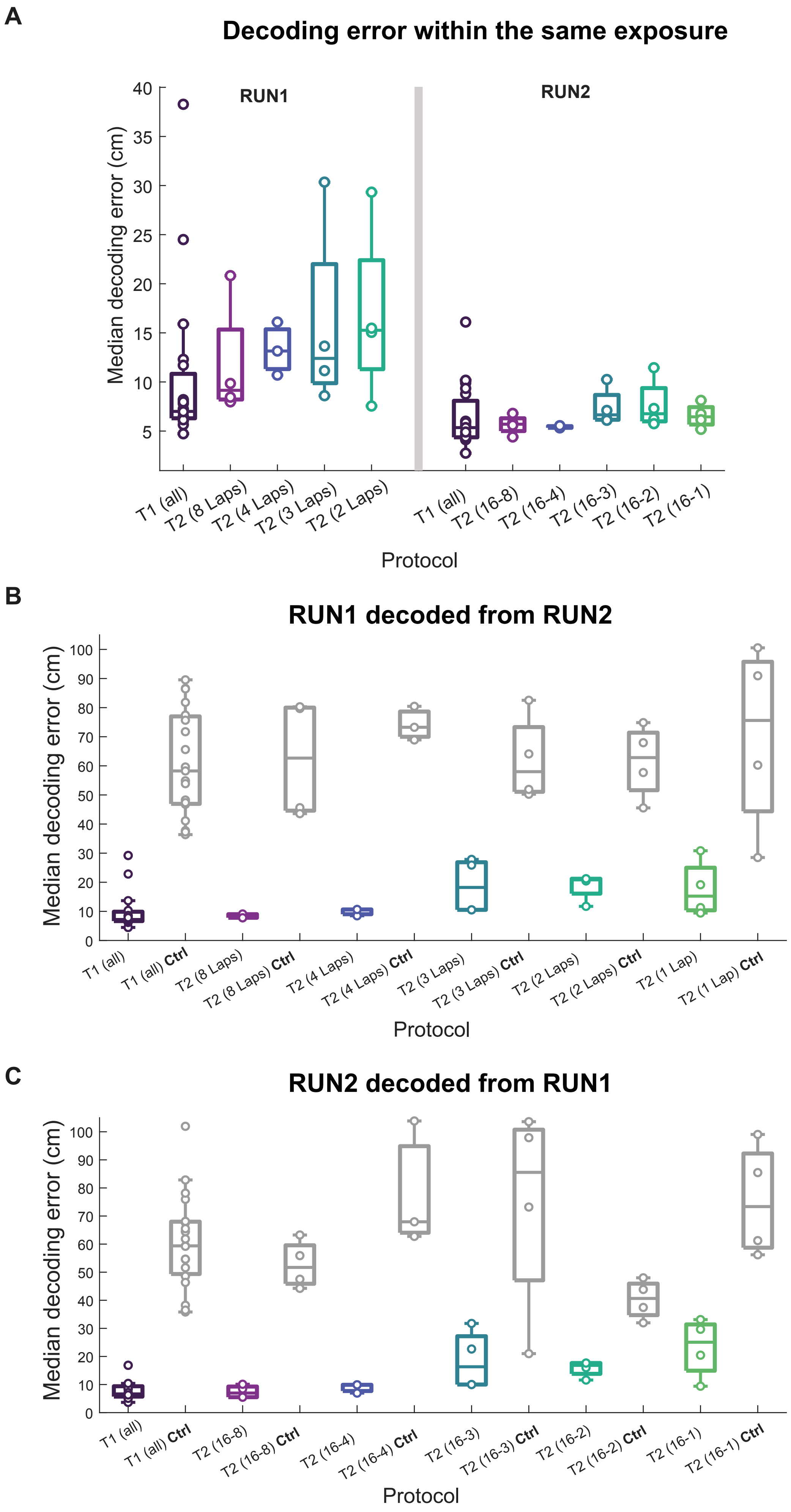


**Fig S2. Summary of median decoding errors across protocols and tracks- (A)** decoding within an exposure based on place fields from the final lap to decode the preceding laps (RUN1 T2) or laps 13-16 to decode laps 1-12 (RUN1 T1 and RUN2 T1+T2), **(B)** RUN1 decoded from RUN2 place fields. **(C)** RUN2 decoded from RUN1 place fields. *Ctrl* **(in B,C)** indicates an control analysis where place fields from the opposite track are used for decoding.

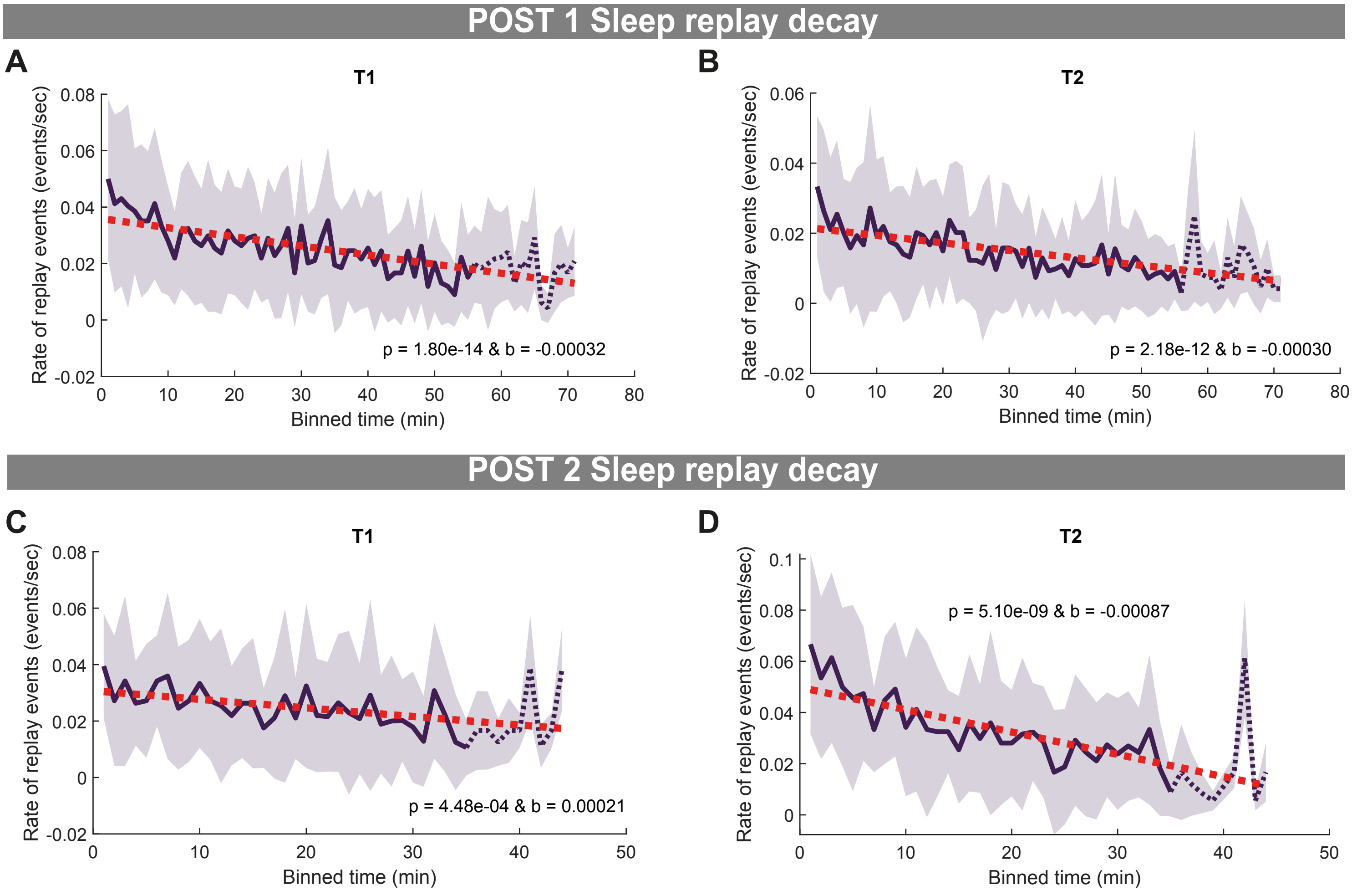


**Fig S3.** Decay of sleep replay rate for track 1 and track 2 events during POST1 and POST2. **(A-D)** Regression between the binned cumulative sleep time and the rate of sleep replay for **(A)** track 1 and **(B)** track 2 events during POST1 and **(C)** track 1 and **(D)** track 2 events during POST2. The shaded region represents the standard deviation across all sessions for a given protocol. The solid line becomes dashed line when less than half of the animals are contributing to the data at each time point.

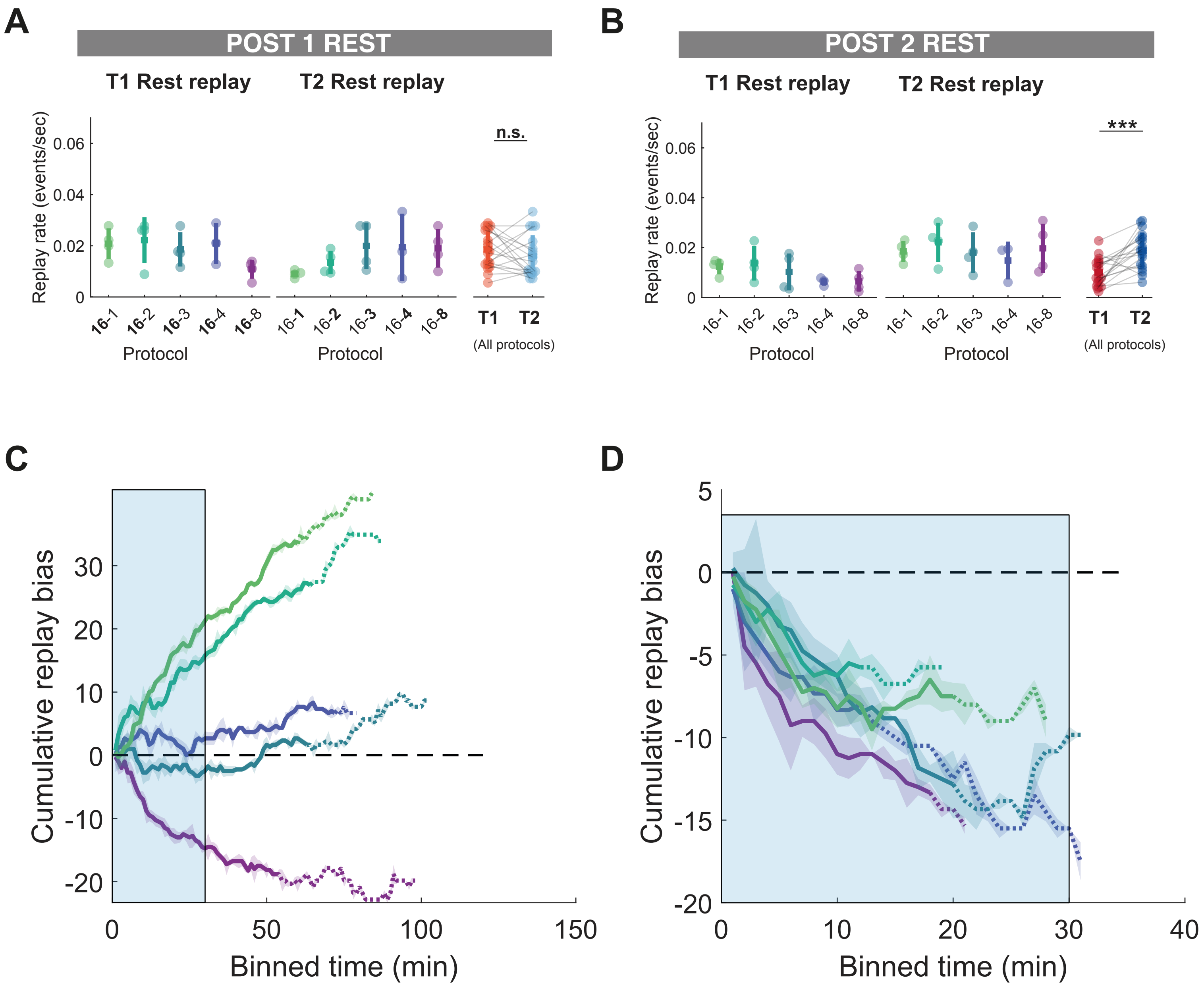


**Figure S4 Hippocampal POST rest replay is sensitive to contextual novelty and familiarity. (A,B)** Rate of sleep replay for track 1 and track 2 during first 30 mins of cumulative sleep of POST1 **(A)** and POST2 **(B)**. **)**. For POST1, the lap number associated with each track during RUN1 was made in bold. For POST2, the lap number was not highlighted as the time spent on two tracks were nearly identical. Each data point represents the mean sleep replay rate within a session, color-coded according to the experimental protocol (***p<0.001, two-tailed Wilcoxon signed rank test). **(C,D)** Cumulative sleep replay bias across protocols during POST1 **(C)** and POST2 **(D)**. The shaded region represents the standard deviation across all sessions for a given protocol. The solid line becomes dashed line when less than half of the animals are contributing to the data at each time point. The light blue box outlined the first 30 mins of cumulative sleep time windows used for analysis in **(A,B).**

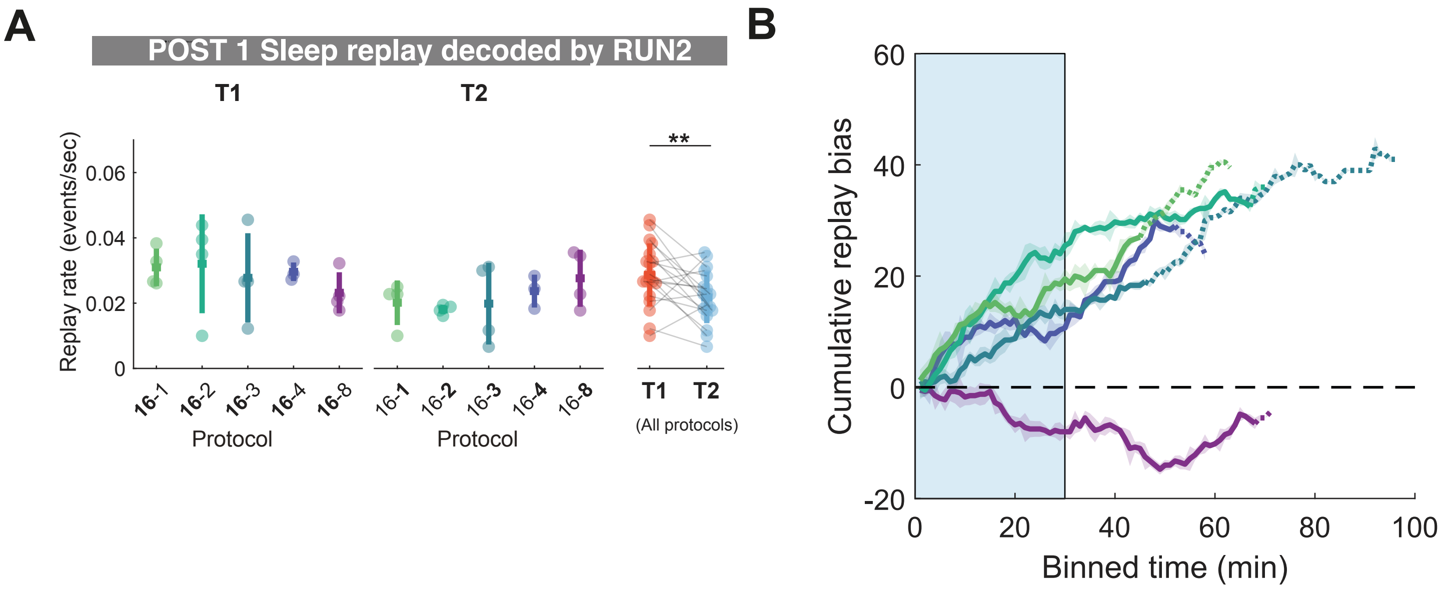


**Figure S5 Sleep replay rate track difference during POST1 remained significantly different when RUN2 place field templates were used for replay decoding.**
**(A)** Rate of sleep replay for track 1 and track 2 during first 30 mins of cumulative sleep of POST1. The lap number associated with each track during RUN1 was made in bold. Each data point represents the mean sleep replay rate within a session, color-coded according to the experimental protocol (***p<0.001, two-tailed Wilcoxon signed rank test) **(B)** Cumulative sleep replay bias across protocols during POST1. The shaded region represents the standard deviation across all sessions for a given protocol. The solid line becomes dashed line when less than half of the animals are contributing to the data at each time point. The light blue box outlined the first 30 mins of cumulative sleep time windows used for analysis in **(A).**

### Awake replay and theta sequence VS POST rest replay

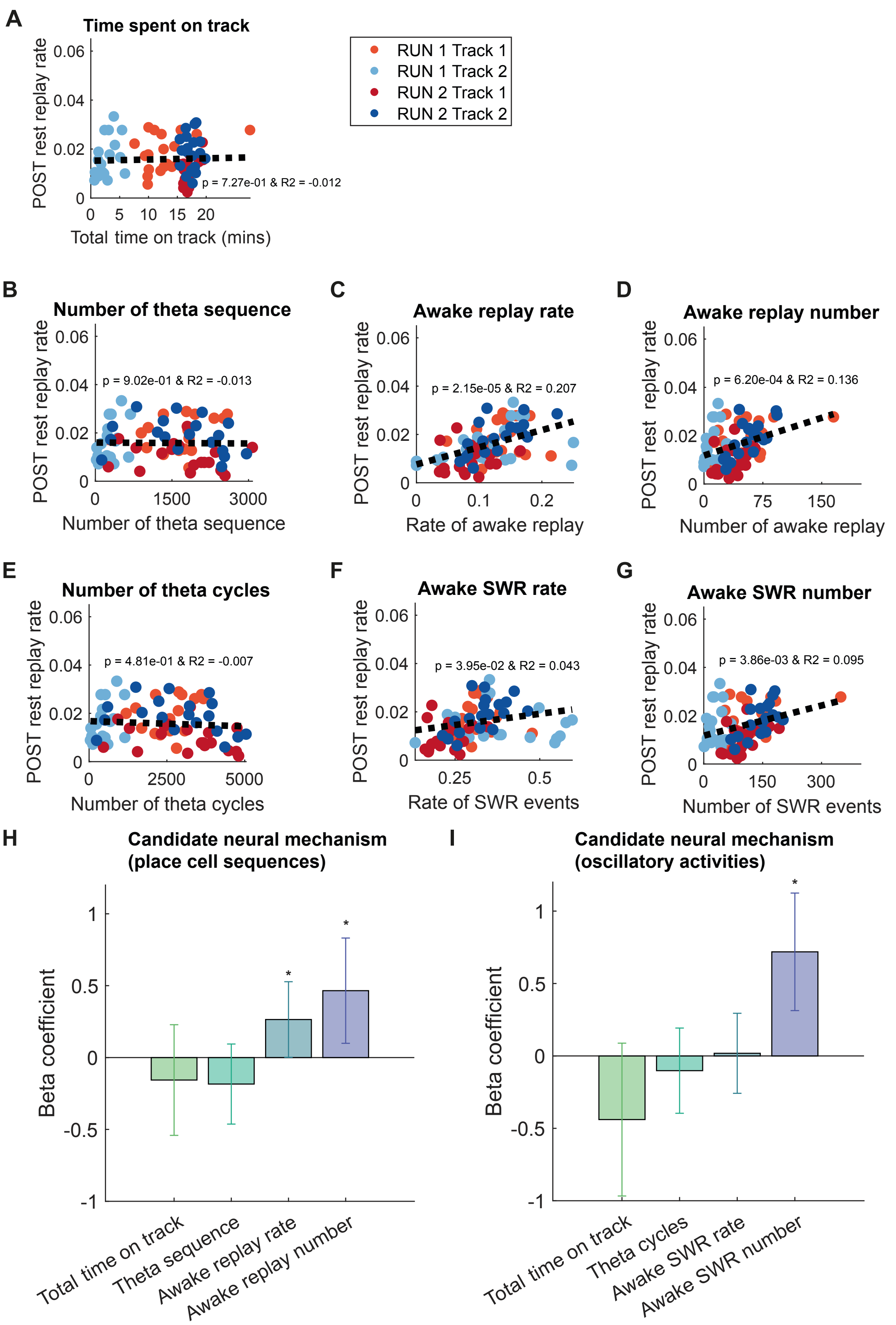


**Figure S6. The predictive relationship between the awake replay and theta sequence and POST rest replay. (A-G)** Simple linear regression of behavioural or neural metric and rate of POST rest replay during first 30 mins of cumulative sleep. **(A)** Time spent on track. **(B)** Number of theta sequence. **(C)** Awake replay rate. **(D)** Awake replay number. **(E)** Number of theta cycles **(F)** Awake SWR rate **(G)** Awake SWR number **(H-I)** Mixed-effect regression for the relationship between candidate neural mechanisms and rest replay **(H)** place cell sequence mechanisms **(I)** Oscillatory events. Asterisks (*) where the 95% confidence interval of the standardized beta coefficient does not overlap with 0.

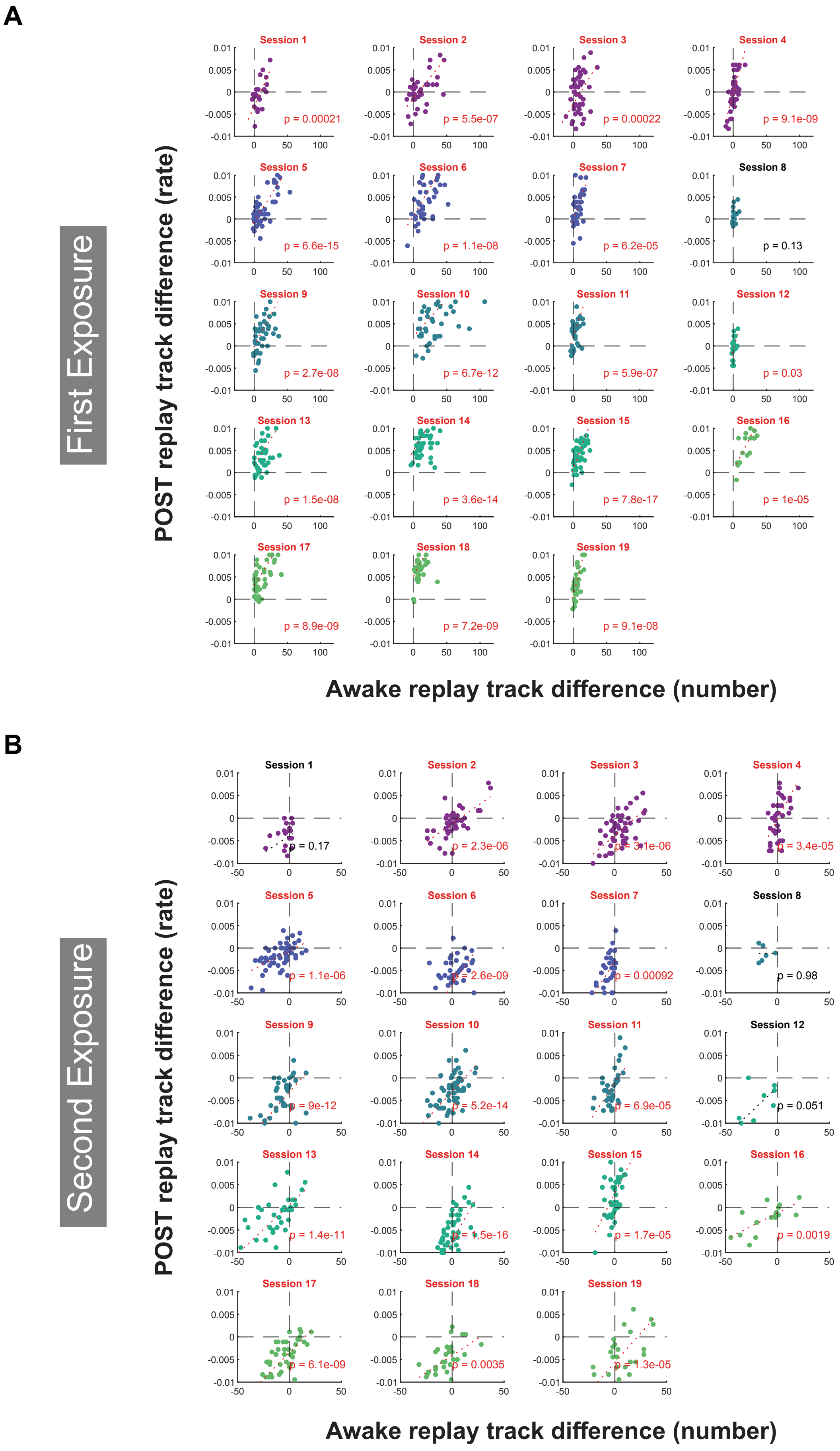


**Figure S7: Place cells participate in more sleep replay events if they participate in more local awake replay events during RUN. (A,B)** Regression between awake replay track difference (number) and POST sleep replay track difference (rate) for first exposure and second exposure. Each data point is a place cell with place fields on both tracks for a given session. For each place cell, the difference in the number of local awake replay events (track 1 - track 2) a given cell was active during RUN was regressed with the observed difference in sleep replay rates (track 1 - track 2) for that cell during the subsequent POST1 **(A)** or POST2 **(B).** Sessions with a statistically significant regression ($p < 0.05$) are highlighted in red.
